# High-throughput submicron-resolution microscopy of entire *C. elegans* populations under strong immobilization by cooling cultivation plates

**DOI:** 10.1101/2021.12.09.471981

**Authors:** Yao L. Wang, Erik L. Jaklitsch, Noa W. F. Grooms, Leilani G. Schulting, Samuel H. Chung

## Abstract

Despite its profound impact on biology, most high-resolution *in vivo* microscopy approaches remain low throughput because current immobilization techniques require significant manual effort. We greatly accelerate imaging of the nematode *Caenorhabditis elegans* by implementing a simple cooling approach to easily immobilize entire populations directly on their cultivation plates. We optimize and characterize cooling immobilization. Counterintuitively, relatively warmer temperatures immobilize animals significantly more effectively than colder temperatures utilized in prior studies. This enhanced immobilization enables clear submicron-resolution fluorescence imaging, which is challenging to achieve with most current immobilization techniques. We demonstrate 64× magnification 3D imaging and timelapse recording of neurons in adults and embryos without motion blur. Compared to standard azide immobilization, cooling immobilization reduces the animal preparation and recovery time by >98%, significantly increasing experimental speed. By obviating individual animal manipulation, our approach could also empower automated imaging of large *C. elegans* populations within standard experimental setups and workflows.

## Introduction

The nematode *Caenorhabditis elegans* is a widely utilized model organism for studying many biological processes. Similar to microorganisms, *C. elegans* are typically cultivated on agarose plates, and they reproduce rapidly by self-fertilization, making them well-suited for studies that require a large sample size. Their transparency and a wide array of labelling techniques allows straightforward visualization of their internal anatomy. Many studies in *C. elegans* image subcellular structures at submicron resolution, which necessitates animal immobilization to prevent image blur. Proper immobilization is especially crucial for techniques involving multiple images in space or time, such as 3D image stacks (*i*.*e*., z-stacks) and timelapse imaging. Typically, immobilization involves manual manipulation of individual animals and mounting them on slides. These time- and labor-intensive procedures restrict experiments to small sample sizes.

Many strategies have been developed to immobilize *C. elegans*, but there are distinct drawbacks for each (see Tab. S1). In brief, chemical methods utilize agents such as sodium azide or levamisole to immobilize by disrupting animal physiology (Reeve, 1988). Mechanical methods, such as polystyrene beads and hydrogel mixtures, physically restrain animals by pressure, viscosity, or friction (Kim, *et al*., 2013). Microfluidic and on-chip technologies constrain animals by pressure, compression, or suction against a membrane or fluid (Guo, *et al*., 2008; Zeng, *et al*., 2008). Lastly, extreme temperatures can also immobilize animals, and most techniques cool animals in flooded wells or microfluidics (Rohde and Yanik, 2011). Despite the many advances in novel immobilization strategies, several significant drawbacks remain: Some of the above strategies require fabrication of slides and significant manual pre- and postprocessing of individual animals. They may increase physiological or physical stress on animals (Manjarrez and Mailler, 2020). Some strategies require complex fabrication and setups or require significant user training. Several methods may also allow small animal movements that blur images, particularly in the nose and pharynx. Notably, none of the current strategies allow for direct imaging on cultivation plates, necessitating time-consuming animal transfer.

Here we address these shortcomings by developing an approach that readily and reversibly immobilizes entire animal populations with minimal user effort. We describe a thermoelectric cooling stage and associated methods that can immobilize *C. elegans* en masse directly on cultivation plates for high-resolution imaging. Our cooling instrument (which we named Copli: <u>co</u>ld <u>pl</u>ate immobilization) stably holds the entire cultivation plate at a desired temperature. We characterize animal movement at several target temperatures and under different cooling rates. Surprisingly, we found that cooling to relatively higher temperatures, specifically 6 °C, minimizes animal movement, though previously reported cooling strategies utilized temperatures between 2-4 °C (Chung, *et al*., 2008; Rohde and Yanik, 2011). Animal movement while cooled consists of bouts of submicron nose tip movement occurring infrequently enough to permit high-resolution brightfield imaging during timescales on the order of minutes. Cooling for an hour does not impact *C. elegans* fecundity or lifespan. Using our novel immobilization method, we obtain submicron-resolution z-stacks and timelapse images of fluorescent neurons in *C. elegans* adults and embryos without blurring from animal motion. We demonstrate that our approach eliminates the need for most of the animal processing steps and greatly accelerates the pace of high-resolution imaging in *C. elegans*. We envision that it could form the basis for automated imaging of large *C. elegans* populations on common experimental setups and workflows.

## Results

### Cooling stage apparatus and temperature distribution

Our instrument design goal is to uniformly cool an entire 55-mm diameter agar cultivation plate to a target temperature, immobilizing animals for high-resolution fluorescence imaging while also preserving transillumination and access to animals. Agar plates are commonly used for cultivating, observing, and manipulating *C. elegans*, so our instrument has broad applicability. As shown in Fig. 1ab, the Copli instrument comprises a thin stage made of an 80-mm diameter sapphire window, a highly conductive copper body, a thermoelectric Peltier heat pump, and a closed-loop liquid cooler. The sapphire window is transparent for transillumination and has a high thermal conductivity. We designed notch cut-outs on the colder side of the sapphire window to improve temperature uniformity across the window. The Peltier is a thermoelectric, solid-state device that pumps heat between its two sides depending on applied voltage.

**Figure 1.**
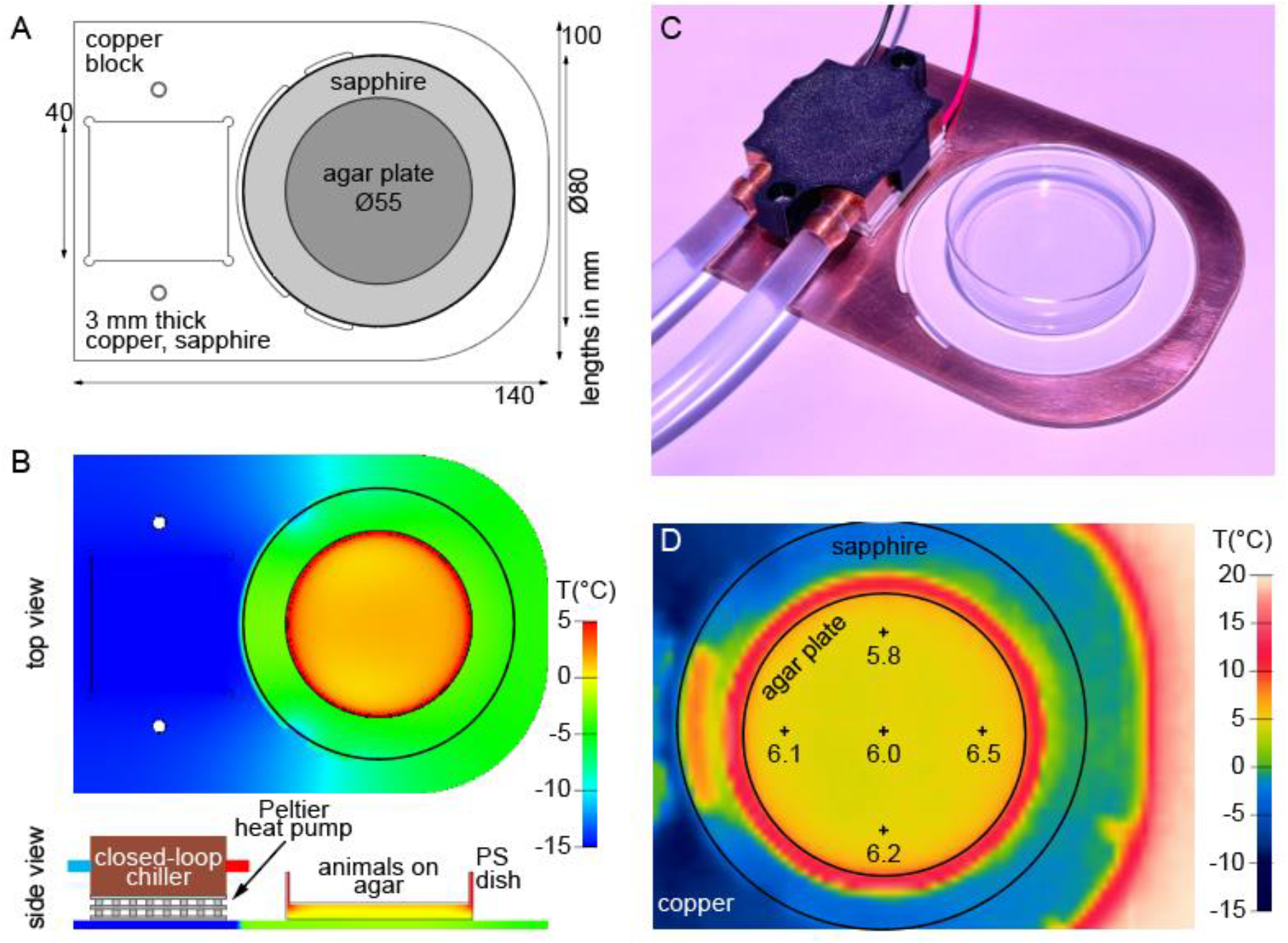
Cooling stage apparatus. (a) Top-down view of cooling immobilization stage. Agar cultivation plates placed on the sapphire window. (b) Thermal modeling of stage and agar plate. Peltier heat pump and liquid cooling system secured to the copper base. (c) Photograph of cooling stage and agar plate. (d) Thermal image of unlidded agar plate and stage cooled to T_set_ = 6 °C obtained by infrared camera. Temperatures at indicated points.

The stage rapidly cools agar plates resting on the sapphire window. Heat leaves the plate to the sapphire window and then transfers to the copper body and Peltier. The Peltier pumps heat to the closed-loop cooler for removal. Thermal simulation of our design predicted a uniform thermal distribution (< 1°C temperature difference from target temperature, T_set_) across the central 40 mm-diameter area of the cultivation plate (Fig. 1b thermal simulation top view). We constructed, assembled, and preliminarily tested the device by achieving and holding a variety of temperatures on the agar surface (Fig. 1c). We measured the thermal distribution of the agar plate and cooling stage, demonstrating an even distribution across the plate’s central area and ensuring that the animals are held at the same temperature for uniform immobilization (Fig. 1d). Thus, Copli’s features include optical transparency for transillumination, high thermal conductivity for cooling efficiency, and uniform temperature distribution across the agar surface.

### General observations of cooling immobilization on agarose plates

Because our study is the first, to our knowledge, to immobilize animals on agar plates by cooling, we began by carefully characterizing the impact of temperature on animal movement. At temperatures ≥ 8 °C, most animals are not immobilized and crawl out of the imaging field of view within minutes. Similar to prior studies of animals in liquid solution, cooling to 1-7 °C slows *C. elegans* movement and stops crawling on plates immediately. Under cooling immobilization, the nose of the animal moves the most of any body part. Pharyngeal pumping is strongly suppressed, the tail also shows some movement, and the body moves the least. Movie S1 shows typical examples of immobilized animal movement on a cultivation plate at 6 °C. The video rate is sped up 50 × to show a 1200-s timespan in a short video. The body parts are immobilized to slightly different levels, consistent with the independent control of these parts (Alkema, *et al*., 2005) and our experience with sodium azide anesthesia (Chung, *et al*., 2013). Because the nose tip exhibits the most movement and contains our target neuron, we verify our cooling immobilization by measuring movement of the nose tip throughout our work described below.

### Optimal cooling temperature for immobilization

We next determined the optimal target temperature, T_set_, to immobilize animals. Previous reports utilized target temperatures of 2-4 °C (Chung, *et al*., 2008; Rohde and Yanik, 2011) but presented minimal quantitative or qualitative characterization of movement. In our experience, some animal movement, however slow or small, persists at all temperatures. We have even observed movement of animals in supercooled buffer held well below 0 °C. By cooling plates to 1-7 °C using our instrument, we investigated animal immobilization in response to a wide temperature range. Surprisingly, we found that a relatively higher temperature, 6 °C, is the optimal temperature for immobilization. Figure 2 shows average nose tip velocity for each minute at specified times after reaching T_set_. At 1-3 °C, nose movement velocity is slow initially but does not greatly decrease over the hour after reaching T_set_ (Fig. 2a). We confirmed that this persistent velocity trend continued for at least 2 h after reaching T_set_ (see below for details). At these velocities (0.5-1.0 μm/s), we could not obtain clear high-resolution images, even with our advanced microscope and sensitive camera. 4 °C represents a transition temperature, where velocity is high initially but decreases to speeds comparable to those at 1-3 °C. At 5 -7 °C, velocity is high initially but, decreases to very low levels after ∼50 min at T_set_. This immobilization appears to take effect progressively over time, in contrast to the immediate immobilization at lower temperatures and earlier times. The velocities at 50-60 min are significantly (*p* << 0.001) below those at 1-3 °C and near or below our resolution limit. As a result, this minimal velocity (< 0.5 μm/s) permits clear submicron-resolution microscopy (see below). Because the final velocities are similar between 5-7 °C, we selected 6 °C as the T_set_ for further study.

**Figure 2.**
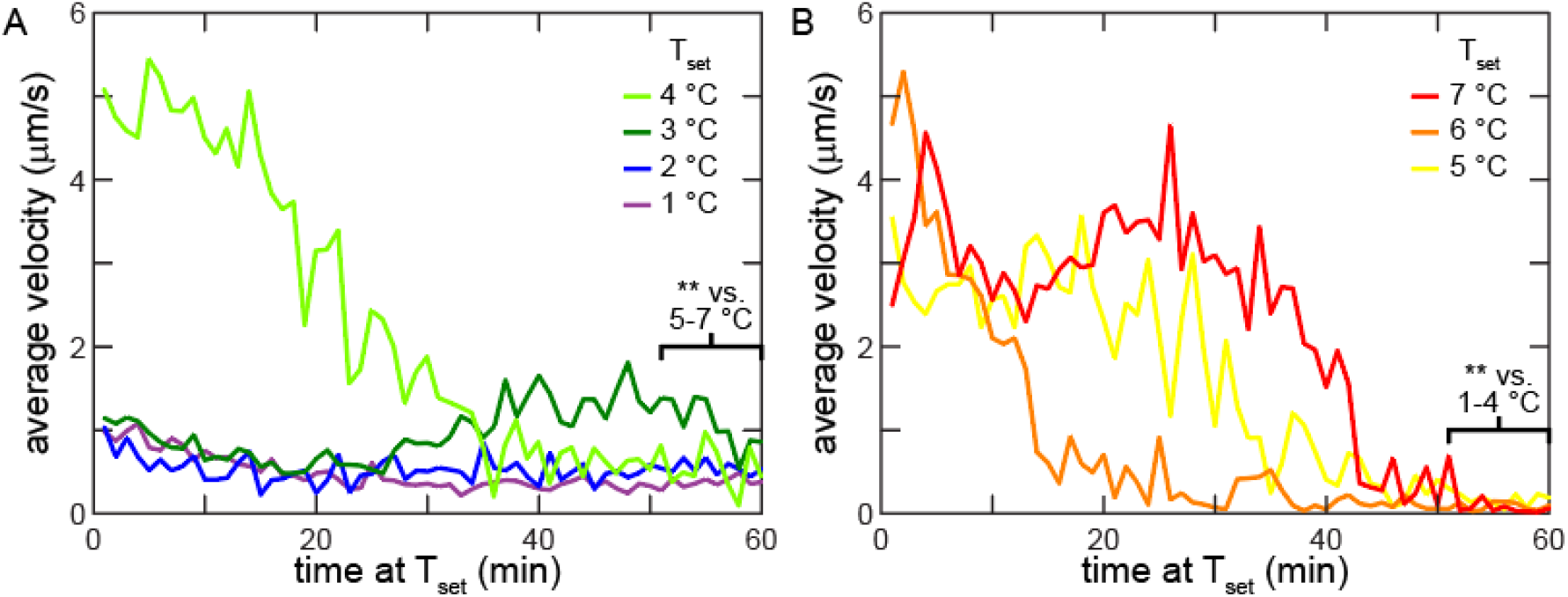
Relatively higher temperatures immobilize animals more effectively. Average nose tip velocity each minute for (a) low temperature (1-4 °C) and (b) high temperature (5 -7 °C) under fast cooling immobilization. *n* = 6 animals. *p* < 0.001 (**) comparing final 10 min of movement for 1-4 °C and 5 -7 °C.

### Different cooling rates produce similar immobilization

We next examined the impact of the cooling rate on immobilization by testing three methods to cool animals down to 6 °C at different rates. We term these methods “slow”, “fast”, and “abrupt”. Figure S1 shows plate temperature profiles for slow and fast cooling methods. Slow cooling uses a typical 4 °C refrigerator to cool down a lidded cultivation plate to 6 °C in about 50 -60 mins. After this first hour in the refrigerator, we transfer this plate to the cooling stage that is pre-cooled, to maintain animals at 6 °C. In fast cooling, we place a plate directly onto the cooling stage to bring the plate from 20 °C to 6 °C. The cooling stage can maintain colder temperatures than the refrigerator; thus, it can cool down animals to 6 °C within 15 mins. Abrupt cooling requires more user effort: we pick animals from their original 20 °C plate directly to a pre -cooled 6 °C plate. Based on strong thermal conductivity and small animal size, we expect animals are cooled to 6 °C in seconds.

As previously stated, immobilization at higher T_set_ is strongest starting ∼50 min after beginning of cooling, and the first hour of slow cooling is in the refrigerator. Thus, we measured velocity for an extended length of time, beginning when the animals reached T_set_ and up to 2 h from the start of cooling (Fig. 3). We compared abrupt, fast, and slow cooling to 6 °C and fast cooling to 1 °C. After cooling to 6 °C, we observe no significant differences in velocity during the 80-120 min range between slow and fast cooling (*p* = 0.08) or between slow and abrupt cooling (*p* = 0.32), indicating that the cooling rate does not require careful tuning to optimize immobilization. By contrast, in agreement with our prior results (Fig. 2), fast cooling to 1 °C allows significantly more movement during the 80 -120 min range than fast cooling to 6 °C (*p* < 0.001). Of the three cooling methods, slow cooling involves the least amount of user time and effort (details in Materials and methods). While certain experiments may require fast or abrupt cooling, we favor the slow cooling method and utilize it for the remainder of this study.

**Figure 3.**
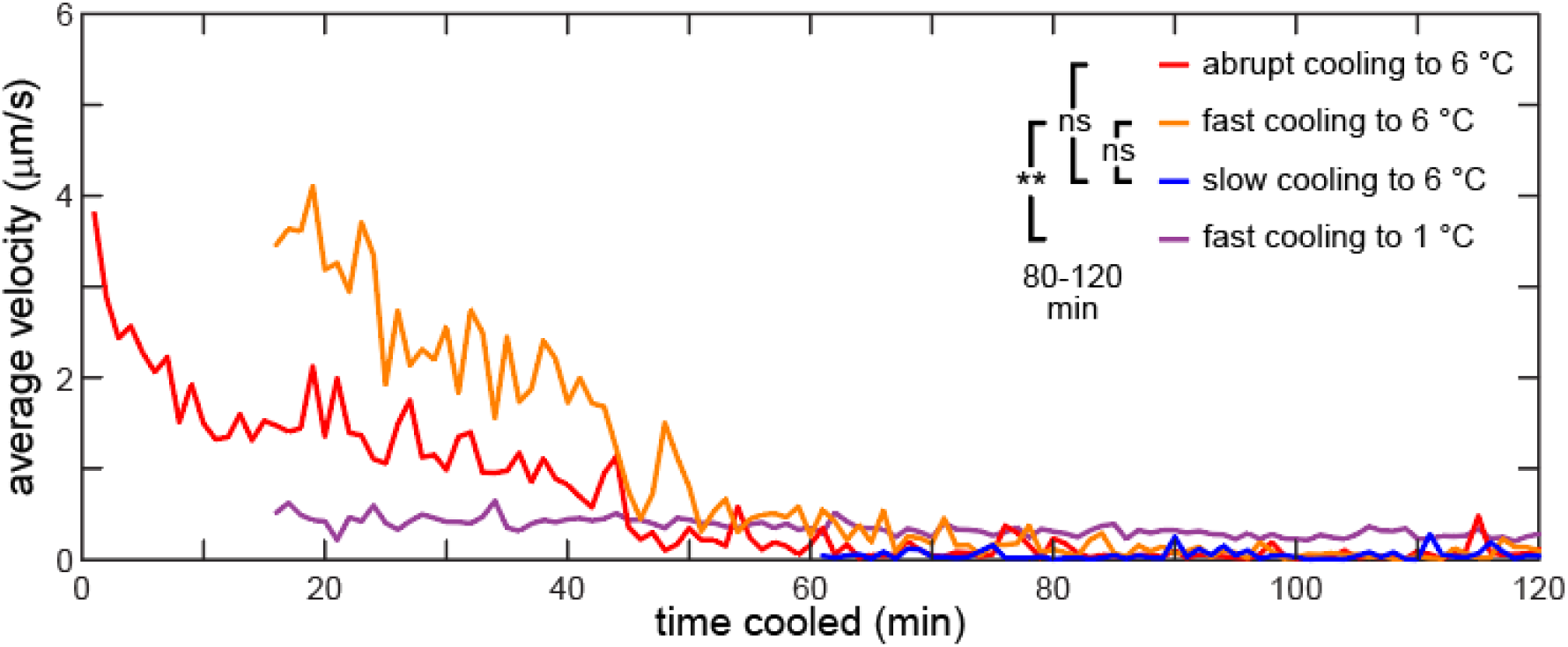
Different cooling rates to 6 °C eventually immobilize similarly. Average velocity of nose tip movement for abrupt, fast, and slow cooling methods to 6 °C and fast cooling to 1 °C. Note tha t fast and slow cooling velocity measurement begins after cooling initiation to allow animals to reach T_set_. *n* = 6 animals for each condition. *p* < 0.001 (**) and not significant (ns) comparisons of movement during 80-120 min.

### Intermittent nose movements under cooling

The goal of our device development is to strongly immobilize animals for imaging, especially for high-resolution microscopy. As such, we ascertained if animals were sufficiently immobilized for imaging by our cooling stage. We measured nose movements each second to more precisely quantify immobilization behavior. Figure 4a shows second-by-second measurements of typical nose movement under 6 °C slow cooling. Similar to sleep and lethargus (Iwanir, *et al*., 2013; Raizen, *et al*., 2008), nose movement under cooling conditions is intermittent, mostly comprised of seconds-long “bouts” of increased activity separated by “windows” with little to no movement. We characterized the distance moved during bouts and the length of the windows. Only 1.4% of movements surpass 1 μm, and 95% of measurements yielded movement below the resolution limit (0.5 μm) (Fig. 4b). Movements below 1 μm do not produce visible blur even at the highest magnification of our microscope (64×) because the motion is largely confined to the tip. Also, most of the experiment duration is contained within longer windows: 96% of the duration is part of windows that are longer than 0.5 min, and 90% of the time is part of windows that are longer than 1 min (Fig. 4c). This level of movement is sufficiently low to image by eye and is also amenable to high-resolution, blur-free brightfield imaging by a camera, which can be performed at most timepoints. Moreover, technology improvements have produced increasingly brighter fluorescent labels, enhanced optics, and efficient cameras. If these tools allow rapid and more sensitive image acquisition, the immobilization requirements may be relaxed. For instance, partial immobilization, achievable at earlier times at 6 °C or all times at 1-4 °C, may be sufficient for imaging with improved technologies, allowing high-speed repetitive immobilizations for longitudinal imaging. In summary, our cooling immobilization approach reduces animal movement sufficiently to allow clear imaging by eye and by camera.

**Figure 4.**
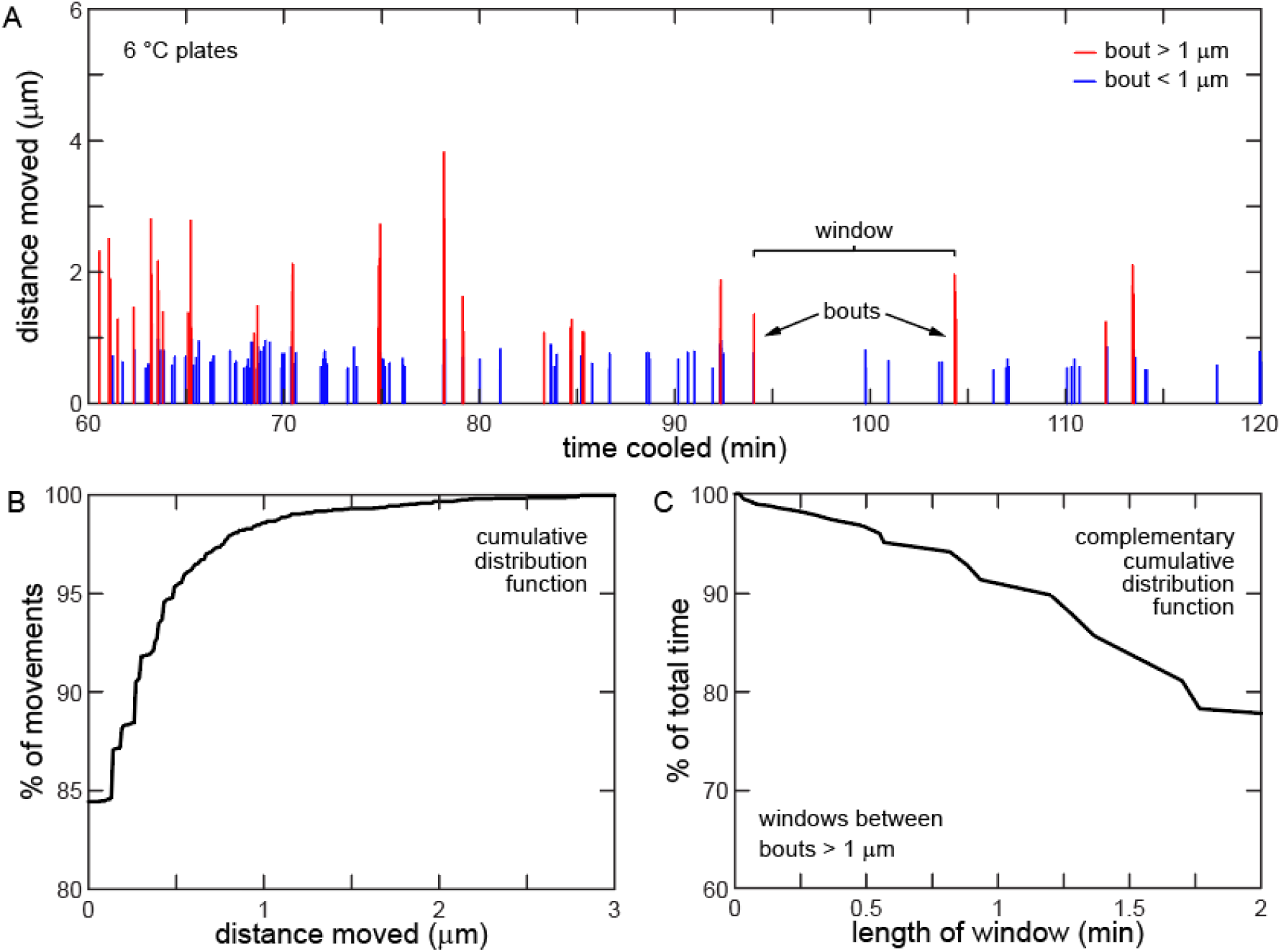
Movement under cooling comprises intermittent bouts with long interstitial windows. (a) Movement measured each second reveals intermittent bouts of increased activity. Movements of < 1 μm do not produce significant image blur. (b) Cumulative distribution function of data in part A. Most movements are small: 99% of movements are < 1 μm. (c) Complementary cumulative distribution function of data in part A. Longer windows comprise most of experiment duration: 96% of duration is part of windows > 0.5 min long.

### Touch stimulus temporarily reactivates movement

We next investigated the impact of touch stimulus on movement. Gentle and harsh touch to *C. elegans* alters the existing behavior of the animal and provokes an escape response (Alkema, *et al*., 2005; Chalfie, *et al*., 2014; Chalfie and Sulston, 1981). Likewise, touching animals in lethargus or sleep also reactivates them (Iwanir, *et al*., 2013; Raizen, *et al*., 2008). We immobilized animals using the slow cooling method to 6 °C and stimulated animals by moving them with a pick at 90 min (*i*.*e*., harsh touch). As expected, touch stimulus reactivates immobilized animals (Fig. S2). Compared to the 10 min before touch stimulus, velocity for the following 10 min is modestly elevated (*p* = 0.006) before returning to the immobilization baseline during the subsequent 10 min. Thus, if animals are disturbed while immobilized, they may require time before returning to full immobilization.

### Animals persist in immobilized state after temperature rises

After a typical experiment involving cooling immobilization, we move plates back to their incubator to return animals to their cultivation temperature. We observed the resumption of movement during warming by turning off the cooling stage. Because animals become strongly immobilized when cooled below 7 °C, we expected that movement would return at a similar temperature when warming animals to their cultivation temperature, 20 °C. Intriguingly, following slow cooling immobilization and maintenance at 6 °C for 1 h, no animals resumed crawling before the plate temperature reached ∼ 10 °C. Even after the plate temperature reached 20 °C, at least 50% of animals remained immobilized until stimulated by gentle touch, whether externally applied or from contact with other animals moving on the plate. When stimulated by touch, animals transition within a few seconds directly from immobilized to crawling that appears normal by eye. Thus, while animals resume apparently normal behavior following cooling immobilization, the transition from immobilized to crawling may require additional study to fully clarify.

### Cooling immobilization does not reduce lifespan or fecundity

Prior studies involving cooling of *C. elegans* indicate that cooling for shorter periods of time does not appear to impact the animal health (Rohde and Yanik, 2011), but longer periods (< 4 h) cause an immediate or progressive decline in health (Ohta, *et al*., 2014; Robinson and Powell, 2016). We assayed viability and fecundity of animals immobilized by cooling for 1 h. First, we immobilized animals at 6 °C (slow cooling) and quantified their lifespan and number of offspring (Fig. S3). Lifespan and fecundity for control and cooled animals are not significantly different (black, blue data). Second, we immobilized animals at 1 °C (fast cooling) to test a harsher immobilization condition. Again, lifespan and fecundity for control and cooled animals are not significantly different (light gray, purple data). We conclude that cooling immobilization up to 1 h does not significantly affect animal health.

### Timelapse imaging and 3D image stack

Illuminating animals with high intensity light stimulates movement in some animals. We describe our observations in the Discussion section. With our setup and animal strains, we note increased movement primarily when the illumination is focused to a size less than the animal length. Under illumination intensities sufficient for fluorescence imaging, most animals remain motionless for a period of tens of second to minutes, allowing clear high-resolution z-stack imaging throughout the animal.

Incompletely immobilized animals produce images with blurred features due to motion during or between images. Blurring must be minimized or eliminated to obtain clear images, particularly for samples with fine features, such as the thin fibers of neurons. By imaging animals immobilized on our cooling stage, we demonstrate high-resolution timelapse and z-stack images of individual neurons without movement-induced blurring. To quantify motion within the imaging window, we obtained timelapse recordings (2 min length, 1 frame/s, 100-ms exposure) of an ASJ neuron expressing green fluorescent protein (GFP). By cross-correlation we tracked the movement of neuron structures within four areas in the field of view that focused on portions of the neuron (Fig. 5a). The distance the neuron structures moved from their original position is shown in Fig. 5b and is usually less than 0.2 μm. Thus, the distance moved during 2 min of imaging is below our resolution limit of 0.5 μm.

**Figure 5.**
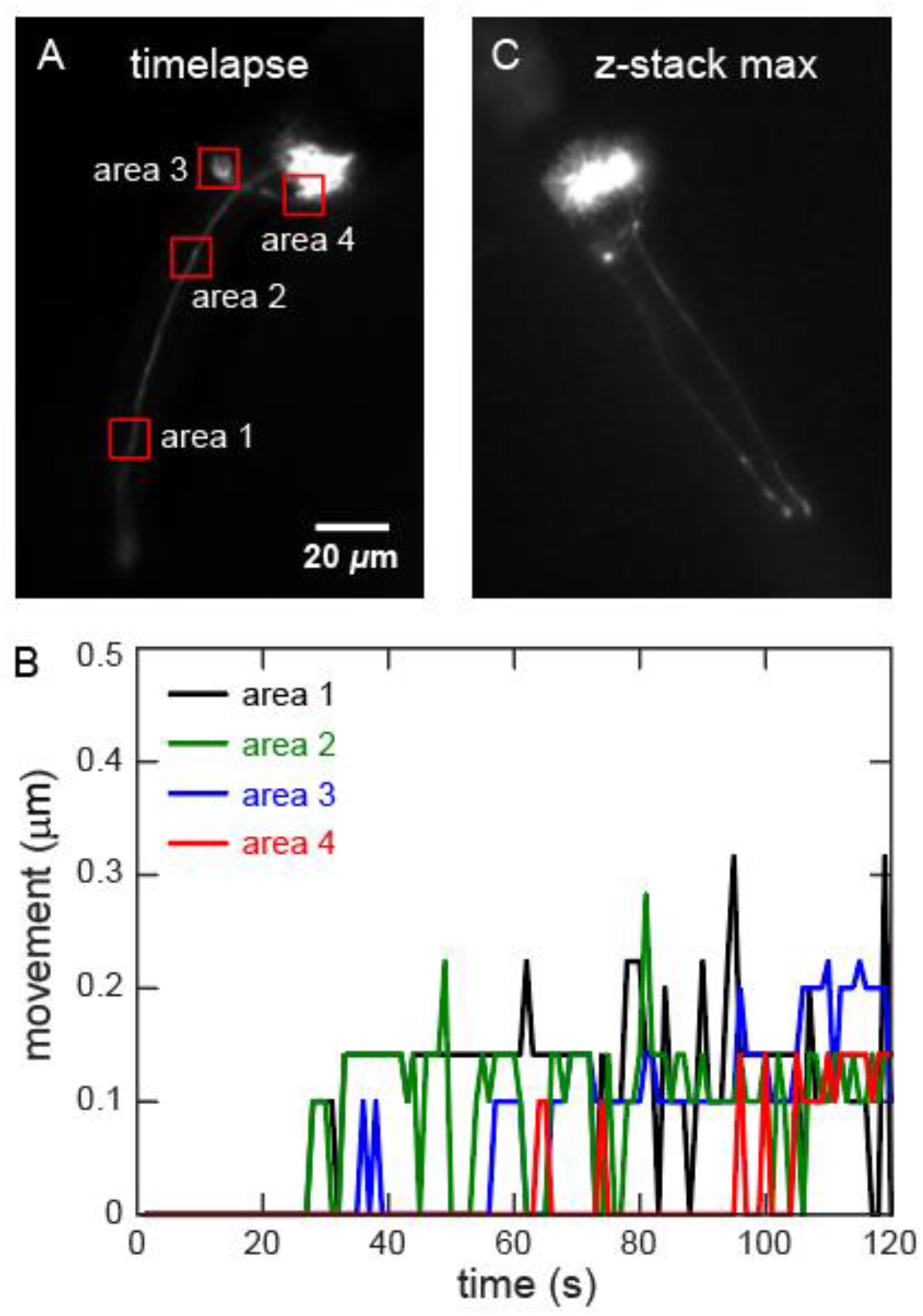
High-resolution imaging under 6 °C cooling immobilization. Fluorescent *C. elegans* ASJ neuron imaged at 64× magnification. (a) Maximum projection of 2 -min timelapse image sequence shows no motion blur. Relative movement of structures in areas 1-4 tracked by cross-correlation. (b) Minimal movement of structures in areas 1-4 during timelapse. (c) Maximum projection of 3D image stack acquired manually. Structures show no motion blur.

We also took a z-stack of the ASJ neuron comprising 41 slices with 1 μm step size taken at 1 slice every 4 s. As shown by the maximum projection of the z-stack in Fig. 5c, while there is haze around neuron structures from inherent out-of-focus and scattered light in widefield imaging (Wang, *et al*., 2021), there is no image blur from motion. Specifically, the neuronal fibers are clearly resolved, even at high resolution. This clear image indicates minimal animal movement during 160 s of imaging, and it underscores the effectiveness of our instrument and approach for immobilization.

### Embryo imaging

Immobilizing *C. elegans* embryos can be challenging due to their protective eggshell. Prior studies immobilized embryos by low temperature (Rabin and Podbilewicz, 2000), mechanical trapping (Cornaglia, *et al*., 2015), and chemical anesthetics (Podbilewicz and Gruenbaum, 2006). While *C. elegans* embryos cannot move any appreciable distance, they rotate and twitch within the eggs. Therefore, we can quantify movement and assess the quality of embryo imaging with and without cooling. We imaged embryos for extended times without cooling and found that animal movement within a 200-ms exposure time blurs cell bodies and precludes clear imaging of small or thin structures, such as neuronal fibers (Fig. 6ac). Embryo cells also move between successive images, so images cannot be aligned or observed with extended-time imaging such as timelapse or z-stacks (Fig. 6bd). We then imaged embryos with slow cooling to 6 °C and found that animal movement was profoundly reduced (Fig. 6eg). There was minimal movement even over timescales > 30 s, allowing clear imaging of neuronal fibers in a single exposure time (Fig. 6fg) and precise alignment of successive images (Fig. 6fh). These timescales are sufficiently long enough to permit acquisition of z-stacks for visualizing 3D structures.

**Figure 6.**
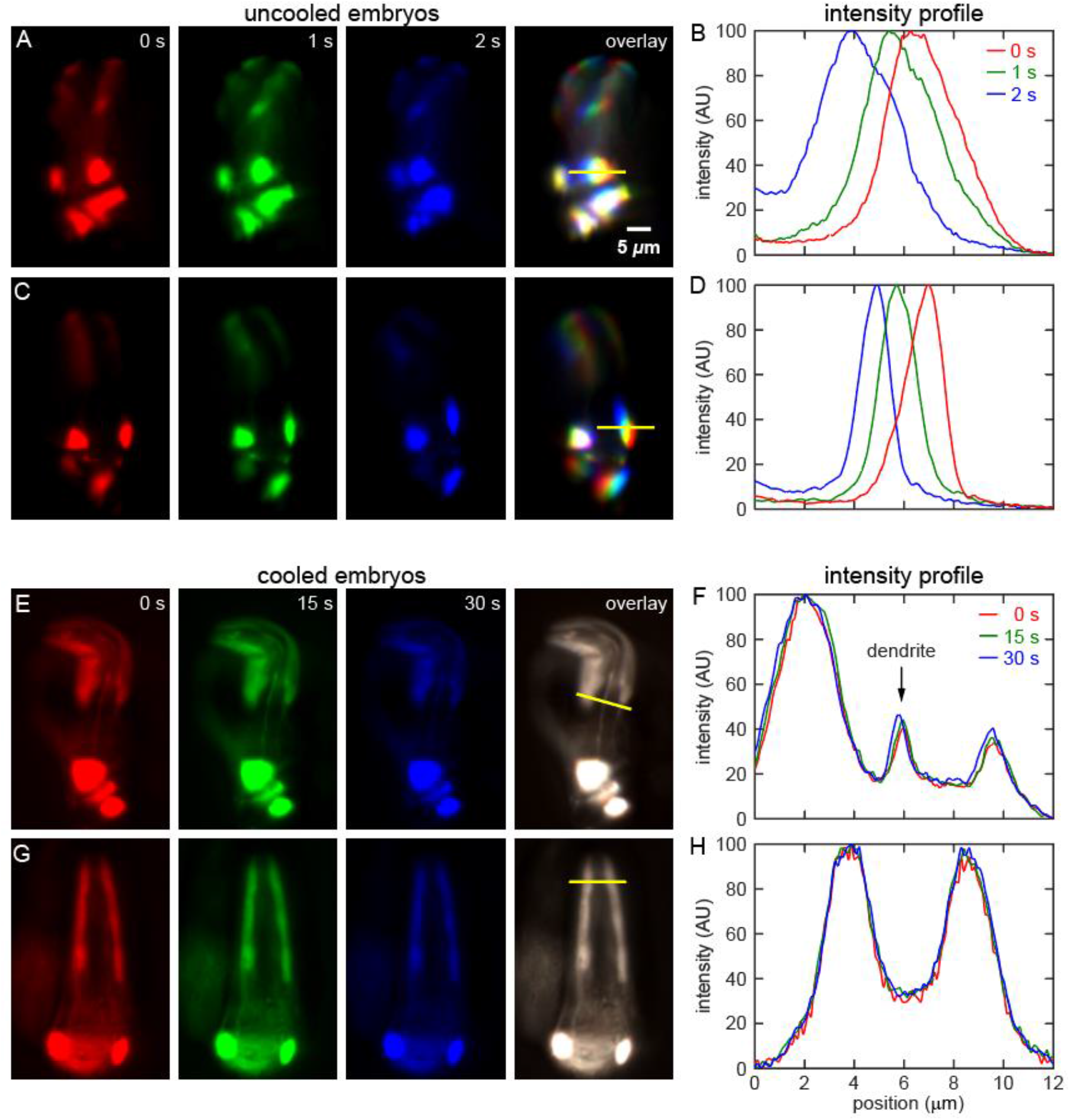
Cooling immobilizes embryo for clear imaging without motion blurring. False-colored fluorescence images of embryo nose (a,c,e,g) and intensity plot profile (b,d,f,h). (a, c) Images taken at 1 s intervals of uncooled embryo. Overlay and profile shows motion blurring. (e, g) Images taken at 15 s intervals of cooled embryo. Overlay and profile indicates that movement is below resolution limit (0.5 μm).

### Cooling immobilization largely eliminates animal processing time

To assess the capacity of our cooling approach to accelerate imaging experiments, we measured time spent imaging 100 animals under standard azide immobilization (Bargmann and Avery, 1995; Chung, *et al*., 2006) and under cooling immobilization. We choose azide immobilization by two reasons. First, azide immobilization is a standard approach and widely used in imaging *C. elegans*. Second, the effectiveness of our cooling immobilization is more similar to azide immobilization than other immobilization techniques.

Azide immobilization requires an involved and time-consuming procedure, which is further extended if animal recovery is required. As shown in Fig. 7a and detailed in Tab. 1, high-resolution single-neuron imaging of 100 animals (selected images in Fig. 7b) requires about 4 h of user effort in total. However, the time actually spent on imaging (Fig. 7a, black bar) is only 1/3^rd^ of the total time. The remainder of the time is spent on pre- and postprocessing (Fig. 7a, red and blue bars), including making agar-azide solution, fabrication of azide slides, mounting and dismounting animals, recovering animals in buffer to remove the anesthetic, and moving animals back to their cultivation plates. Cooling immobilization eliminates nearly all the pre- and postprocessing steps of azide immobilization, reducing the non-imaging time by over 98% (Fig. 7a). Total experimental time is reduced by over 70%, while imaging quality was comparable (selected images in Fig. 7c). As further described in the methods, we believe the timescales measured in our experiment are representative of typical imaging experiments (*e*.*g*., (Chung, *et al*., 2016)). Thus, cooling immobilization greatly accelerates the pace of imaging experiments.

**Table 1.**
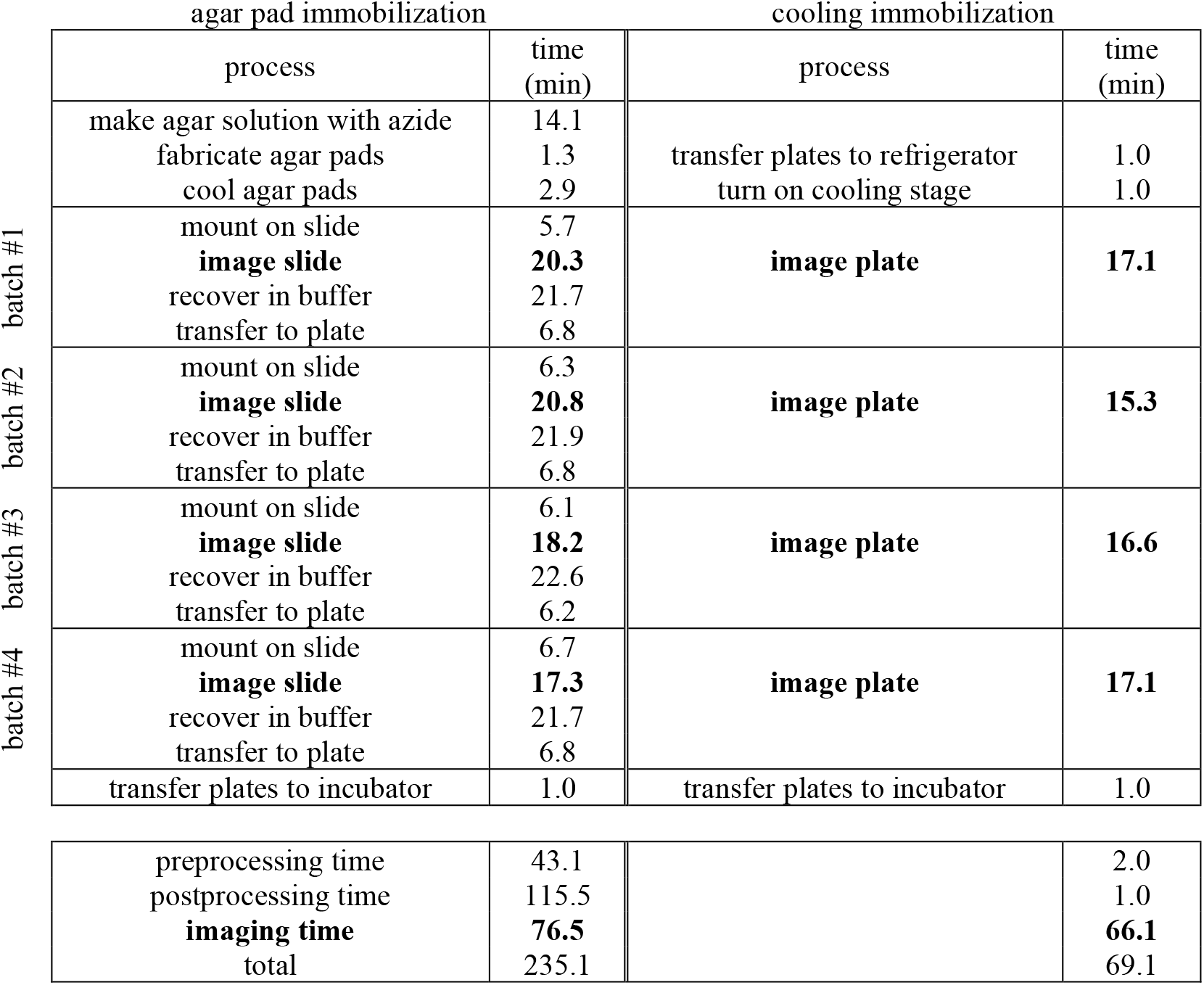
Cooling immobilization eliminates bulk of pre- and postprocessing time. We imaged four batches of 25 animals each for both procedures. Preprocessing includes mounting animals and steps listed prior to the batches. All imaging steps are bolded. Postprocessing time includes recovering animals, transferring animals, and transferring plates.

**Figure 7.**
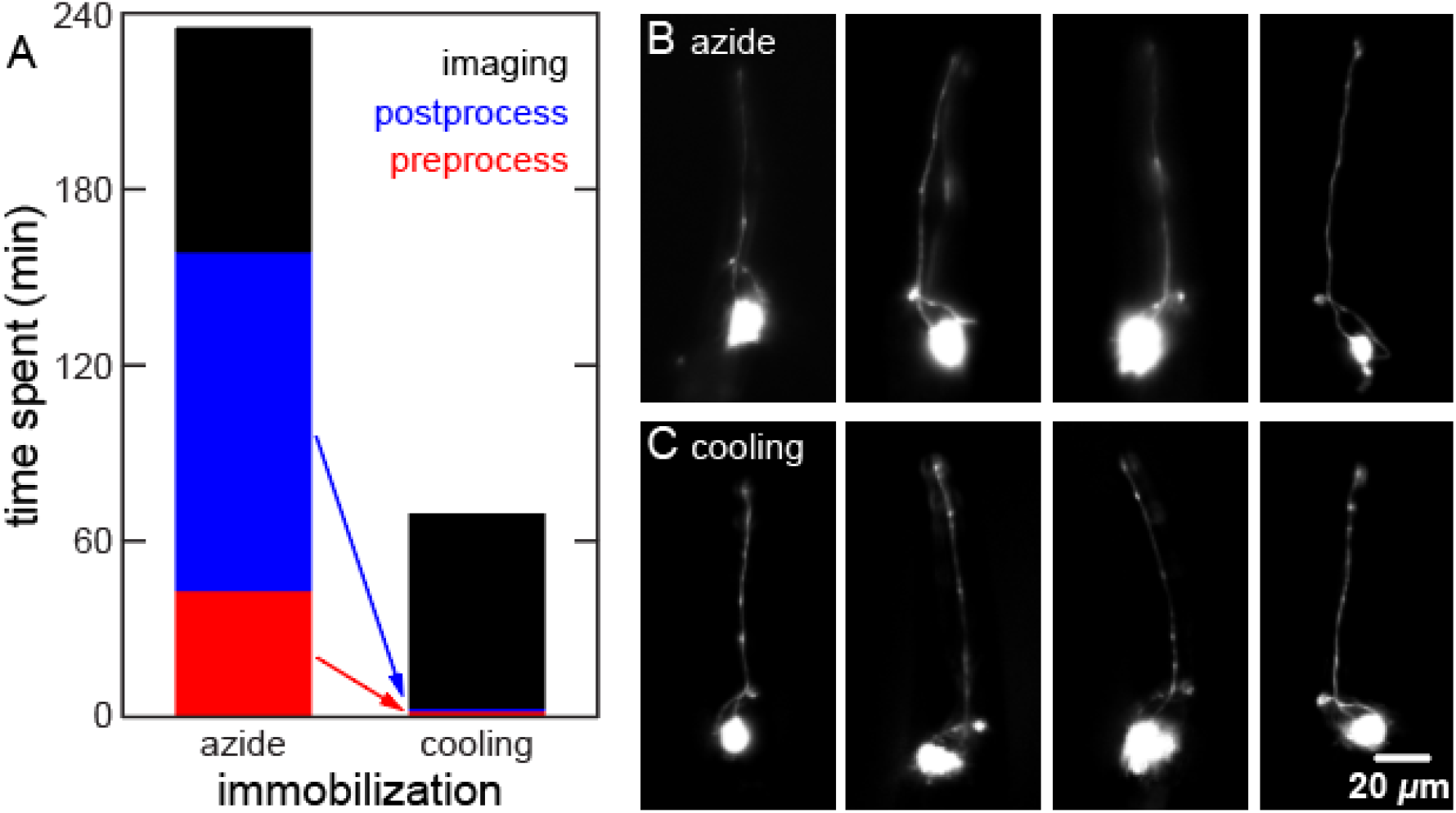
Cooling immobilization reduces non-imaging time by 98%. 100 animals fluorescently imaged under typical azide or cooling immobilization. (a) Time spent on preprocessing, imaging, and postprocessing steps. Preprocessing includes making agar-azide solution, fabricating slides, mounting animals, moving plates. Postprocessing includes recovering animals, removing azide, moving plates. Selected images of ASJ neuron in animal nose under azide (b) and cooling (c) immobilization. Both strategies introduce no motion blur.

## Discussion

### Envisioned workflows

We envision two potential workflows that utilize our cooling approach to enable convenient, reversible immobilization of many animals for high-resolution imaging. In this study, we demonstrated the first workflow, where entire cultivation plates are cooled to 6 °C. Because this method requires ∼50 min to maximally immobilize animals, plates are “precooled” by placing them in a refrigerator for ∼1 h, and subsequent cooling on the stage maintains the immobilization temperature. To maximize efficiency in multi-plate experiments, a plate is processed on the stage while simultaneously precooling the next plate that will be processed. Following imaging of immobilized animals, plates are returned directly to their incubator to bring them back to their cultivation temperature. The second workflow, involving a microchamber device for longitudinal imaging, is further described in the “Future Developments and Applications” section below.

### Advantages of cooling stage and approach

The Copli instrument and approach have several technical capabilities, including some significant improvements over prior immobilization methods. First, large *C. elegans* populations are readily and reversibly immobilized by cooling, allowing observation, imaging, and manipulation of many animals over long time periods. Second, for some applications or genetic backgrounds, immobilization by cooling may be preferable to other methods due to their secondary effects, such as stress response (Manjarrez and Mailler, 2020). For instance, chemical agents such as sodium azide induce stress proteins and thermotolerance in *C. elegans* (Massie, *et al*., 2003). Third, our stage permits high-resolution fluorescence imaging of animals directly on cultivation plates. While *in vivo* microscopy on cultivation plates has been a mainstay technique for decades, it is more often performed by eye and at lower resolution. Our approach enables the study of subcellular processes and cell morphology without manipulating individual animals. Fourth, optical transparency of the sapphire window allows easy observation and imaging by brightfield transillumination, which is a primary method used for many imaging applications in *C. elegans*. Fifth, we designed our cooling stage for straightforward installation into common upright microscope setups. The compact stage is very thin and is easily introduced into and removed from existing setups. Our cooling approach is also amenable to typical *C. elegans* experimental workflows. Finally, most immobilization techniques for high-resolution imaging do not easily allow animal reorientation while imaging. Our cooling stage maintains access to animals, which can be manipulated by a standard platinum pick throughout imaging, at the cost of time to reestablish full immobilization (see “Touch stimulus” section above). These technical capabilities enable broad application of our device and approach to many experiments that require high-resolution *in vivo* microscopy on large numbers of animals.

The Copli instrument and approach also have three practical, user-friendly features. First, our approach requires minimal user effort. Most other reversible immobilization techniques require transfer and manipulation of individual animals to complex setups or slides, which must also be fabricated. Our approach obviates this work by immobilizing animals on their cultivation plate. For high-resolution imaging experiments that involve animal recovery, two-thirds of the experimental time involves non-imaging activities such as animal handling, immobilization, and recovery (Fig. 7a and Tab. 1). Our full sodium azide immobilization procedure (Bargmann and Avery, 1995) requires ∼160 min to process four batches of 25 animals each, not including time spent on imaging animals. Our cooling immobilization procedure requires ∼3 min, representing a savings of >98% of the time expended on non-imaging activities. Thus, our approach greatly accelerates the pace of experiments, enabling the straightforward application of high-resolution imaging to challenging procedures, such as screens. Moreover, because plates hold more animals than slides, cooling immobilization has better economies of scale compared to conventional immobilization. Second, our cooling stage and approach are very simple and straightforward, and they require minimal training to operate. Some immobilization techniques involve complex setups or procedures and thus require significant training and fine-tuning. In contrast, our cooling stage has only a single voltage adjustment to control the immobilization temperature. Thus, our approach’s simplicity makes it intuitive as well as resistant to failure. Finally, our device is accessible and inexpensive: we assembled it from easily attainable and affordable components, totaling a cost of ∼$500. In summary, our device and approach have significant technical capabilities in a very user-friendly design.

### Key technical developments and novel findings

In our study, we develop a novel, compact thermoelectric stage that cools entire cultivation plates to immobilize populations of *C. elegans* for imaging. Our stage cools plates to a uniform temperature distribution across the agar surface that is stable over long time periods. We describe animal movement under many cooling conditions in far greater detail than previously reported. We demonstrate that cooling has minimal impact on lifespan and fecundity. Our cooling stage successfully immobilizes animals sufficiently to allow high-resolution brightfield imaging. Despite some movement arising from high-intensity illumination for high-resolution fluorescence imaging, we demonstrate the capacity of our immobilization approach to perform blur-free time-lapse and z-stack fluorescence imaging. In summary, our cooling stage represents a novel and effective way to reversibly immobilize large populations of *C. elegans* for high-resolution brightfield and fluorescence imaging.

During our study, we also encountered some interesting and unusual animal behavior. First, previous studies immobilized animals between 2-4 °C (Chung, *et al*., 2008; Rohde and Yanik, 2011). By careful and systematic assay, we find that optimal immobilization of *C. elegans* occurs at a counterintuitively higher temperature of 5-7 °C after animals have been cooled for ∼50 min. In contrast, animals cooled to 1-3 °C maintain a relatively stable level of movement that is significantly higher than at 6 °C. Animals held at 1 °C on ly slightly reduce their movement even after 2 h. In fact, a higher immobilization temperature is preferable as it is easier to maintain, has lower risk of freezing animals, and is presumably a more benign environment. Second, animal behavior under cooling and under sleep share some characteristics. Similar to sleep, animals immobilized by cooling spontaneously switch periods of movement and quiescence. After cooling immobilization, many animals persist in the immobilized state even after warmed to cultivation temperatures but resume normal movement rapidly following gentle touch. Further studies are required to determine if cooling immobilization exhibits other hallmarks of sleep, or if it is more akin to partial anesthesia.

### Limitations of cooling stage and approach

There are several limitations of our approach. First, animals raised at 15 °C continue to crawl at our standard immobilization temperature of 6 °C, indicating that cultivation temperature modulates cooling immobilization. This is not unexpected as prior studies have shown that raising *C. elegans* at colder temperatures improves cold tolerance (Murray, *et al*., 2007). Thus, additional fine-tuning of our approach is required if animals are not cultivated at 20 °C. Likewise, we also speculate that some temperature-sensitive mutant strains of *C. elegans* may require adjustment of immobilization temperature, as we have optimized our approach for the wild-type background. Second, the speed of immobilization is slower than most other methods, but we expect that the easy step of pre-cooling plates overcomes this weakness for most applications. Third, imaging fine structures at locations that exhibit relatively more movement, such as the dendritic cilia at the nose tip, could be challenging.

Finally, high-intensity illumination for fluorescence imaging at high-resolution stimulates movement in some animals. Our current investigations are aimed at quantifying and minimizing this effect. By eye, the nose appears to be more sensitive than the tail or the body to light. The location of GFP expression may affect light sensitivity as FLP-fluorescent animals could be illuminated with >10× more light intensity than ASJ-fluorescent animals before stimulating movement. Also, green light illumination (red fluorescence imaging) triggers much less movement than blue light (green fluorescence imaging). These gradations of sensitivity are consistent with prior findings that neurons in the nose, and particularly the ASJ, contribute strongly to wavelength-dependent light sensitivity in *C. elegans* (Ward, *et al*., 2008). Despite this sensitivity, we are still able to demonstrate high-resolution fluorescence imaging at the submicron level without motion blur.

### Future developments and applications

Our cooling approach and platform could be developed in several ways to greatly enhance its capabilities. A microfluidic or microchamber (Bringmann, 2011) device utilized with our cooling approach could permit longitudinal imaging of individual animals. A thin device would enable rapid, repeated cooling for brief imaging windows with minimal impact on animal health (Rohde and Yanik, 2011). Such a device would be useful for tracking development or neuroregeneration following injury. In our study, we also completed procedures manually, but one advantage of our approach is that it is amenable to automation. Commercial plate handlers can move plates to instruments and could be easily adapted to load plates onto our cooling stage. The cooling immobilization and imaging could also be automated through multiple coolers and translation stages. There is also great potential for integrating our approach with image processing and segmentation algorithms to enable automated recognition of cellular components and targeting for laser surgery. Accelerating the imaging by itself, however, would not reduce the substantial amount of time spent on non-imaging activities, highlighting the need to develop immobilization platforms to fully exploit automation. The developments listed above would significantly extend the capabilities and accelerate the throughput of our platform, allowing its application to a wide array of questions that are currently unapproachable.

## Acknowledgements

We thank the members of the Chung Laboratory for feedback on the manuscript. We thank Neil Patel (Bioengineering, Northeastern Univ.) for assistance with animal nose tip tracking. We thank Max Heiman (Genetics, Children’s Hospital Boston) for providing CHB4526 for embryonic imaging. We acknowledge Wenchao Zhu (Mechanical Engineering, Northeastern Univ.) for technical support in thermal distribution measurement. We acknowledge Xin Sun (Bioengineering, Northeastern Univ.) for technical support in 3D printing.

## Author contributions

Several people contributed to the work described in this paper. SHC conceived the idea. YLW, ELJ, and NWFG carried out experiments, and all authors analyzed the data. SHC supervised the experiments and the development of the manuscript. ELJ and SHC wrote the first draft of the manuscript; all authors subsequently took part in the revision process and approve the final copy of the manuscript.

## Declaration of interests

The authors declare no competing interests.

## Multimedia files

**Movie S1. 20-min timelapse recording of immobilized animals**. We acquired a video of three wild type N2 animals immobilized on their cultivation plate under 6 °C, using 10× magnification. We speed up the 20-min long video by 50× to facilitate viewing. Animal movements occur at times (s): Left animal nose tip-99, 182, 392, 822, 1045; tail-359, 723. Center animal tail-133, 297. Right animal nose tip-0, 292, 469, 488.

## STAR Methods

### Device design and components

We utilized Solidworks to design and model the thermal distribution of the cooling stage. Components: Peltier heat pump (CP854705-2, 40 × 40 mm, from CUI Devices, Lake Oswego in Oregon); 99.9% pure copper stage block (100 × 140 × 3 mm, machined); sapphire window (80 mm diameter, 3 mm thickness, from Altos Photonics, Bozeman in Montana); Thermal paste to connect parts (Arctic MX-4, from Florence in South Carolina); liquid CPU Cooler (Hydro Series H100x, from Corsair, Fremont in California); copper liquid cooling heatsink (40 × 40 × 10 mm, replacing the original bulky heatsink, unbranded); custom installation bracket fabricated by FDM (Fused Deposition Modeling) 3D printing of polylactide filament.

### Plate temperature measurements

We measured the thermal distribution of the agar plate using a calibrated thermal camera (Teledyne FLIR C2, Thousand Oaks in California). We embedded a wire digital thermocouple (Proster, San Jose in California) 1 mm into the agarose on plates to confirm camera readings and track agarose temperature over time.

### Nematode handling

We maintained *C. elegans* on Bacto agar plates under standard conditions at 20 °C (Brenner, 1974). We utilized three strains in our study: a wild-type N2 strain, a fluorescent strain with *daf-11(sa195)V; ofIs1[ptrx-1::trx-1::gfp]* for ASJ neuron imaging, and CHB4526 *hmnEx2428[pIM48(egl-13pro::GFP), pRF4]* for embryo BAG and URX neuron imaging. We utilized N2 for all experiments except for fluorescence imaging, where we utilized the strains with green fluorescent protein (GFP). We synchronized large populations of animals for experiments by placing ∼30 adults on plates to lay eggs for 4 h. We transferred animals between plates using a pick. For the touch stimulus experiment, we used a pick to move animals at least 1 cm away from and then back to their original location.

## Cooling methods: slow, fast, and abrupt

In general, the parameters of the cooling methods, such as voltages and times, that lead to the desired performance, will depend on specific parameters of the cultivation plates and stage, such as the amount of agar used to make the plates, the efficiency of the Peltier stage, and the material conductivities.

### Slow cooling

In our study, we performed slow cooling to a target temperature (T_set_) of 6 °C only. We first moved a 20 °C plate with its lid to a 4 °C refrigerator for 1 h. Then, we moved the plate directly to our cooling stage (pre-cooled for 10 min with 4.8 V across the Peltier), with the plate’s lid off for better imaging and worm manipulation.

### Fast cooling

We first moved a 20 °C plate with animals to a cold stage (cooled for 10 min at 12 V), with plate lid off. When the plate temperature reached T_set_ + 5 °C, we turned the voltage to a lower value (4.8 V for T_set_ = 6 °C and 7.1 V for T_set_ = 1 °C) to avoid over-cooling and maintain plates at T_set_. This method cools down plates to different target temperatures in ∼15 mins.

### Abrupt cooling

In our study, we performed abrupt cooling to 6 °C only. We moved animals from their 20 °C cultivation plates directly to a pre-cooled 6 °C plate resting on the cooling stage.

### Tip-Tracking

We first captured videos of animals under the stated conditions at 20 frames per second (FPS). Then, we extracted 1 frame per second from those videos using a custom MATLAB script. We manually tracked the nose tip position in those frames using the Multi-point Tool in ImageJ and calculated velocities for each second in MATLAB. We averaged the individual velocities of 60 s to produce average velocities for each minute.

### Lifespan and fecundity analysis

We collected N2 wild-type animals in the L4 lifestage on two plates. We left the control plate in the cultivation incubator. The cooled plate was processed by the slow cooling to 6 °C and fast cooling to 1 °C methods described above. We held animals at T_set_ for 1 h. We then moved the cooled plate back to the cultivation incubator.

### Lifespan

A few hours after cooling or control treatment, we transferred animals to separate plates to avoid crowding. We observed animals twice daily following established procedure (Larsen, *et al*., 1995) to assess mortality. We transferred animals to new plates when needed to avoid starvation.

### Fecundity

A few hours after cooling or control treatment, we transferred animals individually (*i*.*e*., “singled”) to plates. Three days later, we moved the singled animals to new plates and counted the number of hatched offspring for each animal. We repeated this process until there were no new offspring.

### Fluorescence imaging

We used a Leica M205 FA widefield fluorescence stereomicroscope with 5×, 0.5 numerical aperture (NA) objective, and an Andor Zyla 4.2 sCMOS camera (Oxford Instrument; Concord, MA) to obtain images. We imaged neuronal GFP at 64× magnification, with theoretical 0.5 μm transverse and 4 μm axial resolution. We immobilized animals by slow cooling to 6 °C. We performed single-frame, timelapse, and z-stack fluorescence imaging on the ASJ neuron with 100-ms exposure time. For timelapse imaging, we used 1 FPS for 2 minutes to obtain 120 frames in total. For z-stack imaging, we used a 3-axis stage (Thorlabs MBT616D, Newton in New Jersey) to obtain slices over a 40-μm z range, with 1-μm z step, and 4-s time interval between slices. For embryo imaging on the BAG and URX neurons we used 1 FPS with 200-ms exposure time.

### Measurement of time expenditure for azide and cooling immobilization

To compare the efficiency of sodium azide and cooling immobilization, we recorded time spent on the processes needed to acquire a 2D image of the ASJ neuron in each of 100 animals (Tab. 1). We immobilized, manually located, imaged, and recovered animals in 4 batches, which reduces the length of time the animals are immobilized and improves animal viability. For comparison, we categorized processes into three general procedures: preprocessing, imaging, and postprocessing.

We performed sodium azide immobilization as previously described (Chung, *et al*., 2006). The preprocessing steps include making the agar-azide solution, fabricating and cooling agar pads, and mounting and arranging the animals on the agar pad. The postprocessing steps include recovering animals in buffer to rapidly remove azide, transferring recovered animals to cultivation plates, and moving cultivation plates back to the incubator. We believe the times we recorded accurately represent user time expended in a typical high-resolution imaging experiment with azide immobilization. The time would increase if animals are sorted into groups (*e*.*g*., surgery vs. mock), and the time would decrease if animal recovery is not required or if the agar-azide solution is reused.

We performed slow cooling immobilization to 6 °C as described above. To optimize comparison with azide immobilization, we seeded with OP50 bacteria an area equal in size to our agar pads. Animals were largely confined to this area. We envision our cooling immobilization will be utilized on entire populations directly on their primary cultivation plates, so no transfer of animals is required. The preprocessing steps includes transferring plates to refrigerator and turning on the cooling stage. The postprocessing steps includes transferring plates to incubator.

### Statistics and interpretation of results

For comparing nose tip velocities, we calculated *p*-values using the *t* test: For different temperature settings or different cooling rates, we utilized the paired, two-tailed *t* test. For examining touch stimulus, we utilized the unequal variance, two-tailed *t* test.

For assessing the impact of cooling on viability and fecundity, we calculated *p*-value by using log-rank test and the unpaired, equal variance, two tailed *t* test, respectively. Data are represented as average ± standard deviation (SD). In figures, we indicate values that differ significantly at *p* < 0.05 (*) and *p* < 0.001 (**) levels.

## Notes

### Competing Interest Statement

The authors have declared no competing interest.

### Summary of Updates

We performed new experiments about C. elegans embryos imaging and animal processing time, and add those experiment results to the new manuscript.

## References

Alkema, M.J., Hunter-Ensor, M., Ringstad, N., and Horvitz, H.R. (2005). Tyramine Functions Independently of Octopamine in the Caenorhabditis elegans Nervous System. Neuron 46, 247–260.

Bargmann, C.I., and Avery, L. (1995). Laser killing of cells in Caenorhabditis elegans. Methods Cell Biol 48, 225–250.

Brenner, S. (1974). The genetics of Caenorhabditis elegans. Genetics 77, 71–94.

Bringmann, H. (2011). Agarose hydrogel microcompartments for imaging sleep- and wake-like behavior and nervous system development in Caenorhabditis elegans larvae. J Neurosci Methods 201, 78–88.

Chalfie, M., Hart, A.C., Rankin, C.H., and Goodman, M.B. (2014). Assaying mechanosensation. WormBook : the online review of C elegans biology, 10.1895/wormbook.1891.1172.1891.

Chalfie, M., and Sulston, J. (1981). Developmental genetics of the mechanosensory neurons of Caenorhabditis elegans. Dev Biol 82, 358–370.

Chung, K.H., Crane, M.M., and Lu, H. (2008). Automated on-chip rapid microscopy, phenotyping and sorting of C. elegans. Nat Methods 5, 637–643.

Chung, S.H., Awal, M.R., Shay, J., McLoed, M.M., Mazur, E., and Gabel, C.V. (2016). Novel DLK-independent neuronal regeneration in Caenorhabditis elegans shares links with activity-dependent ectopic outgrowth. Proc Natl Acad Sci USA 113, E2852–E2860.

Chung, S.H., Clark, D.A., Gabel, C.V., Mazur, E., and Samuel, A.D. (2006). The role of the AFD neuron in C. elegans thermotaxis analyzed using femtosecond laser ablation. BMC Neurosci 7, 30.

Chung, S.H., Schmalz, A., Ruiz, R.C.H., Gabel, C.V., and Mazur, E. (2013). Femtosecond Laser Ablation Reveals Antagonistic Sensory and Neuroendocrine Signaling that Underlie C. elegans Behavior and Development. Cell Reports 4, 316–326.

Cornaglia, M., Mouchiroud, L., Marette, A., Narasimhan, S., Lehnert, T., Jovaisaite, V., Auwerx, J., and Gijs, M.A.M. (2015). An automated microfluidic platform for C. elegans embryo arraying, phenotyping, and long-term live imaging. Sci Rep 5, 10192.

Guo, S.X., Bourgeois, F., Chokshi, T., Durr, N.J., Hilliard, M.A., Chronis, N., and Ben-Yakar, A. (2008). Femtosecond laser nanoaxotomy lab-on-a-chip for in vivo nerve regeneration studies. Nat Methods 5, 531–533.

Iwanir, S., Tramm, N., Nagy, S., Wright, C., Ish, D., and Biron, D. (2013). The Microarchitecture of C. elegans Behavior during Lethargus: Homeostatic Bout Dynamics, a Typical Body Posture, and Regulation by a Central Neuron. Sleep 36, 385–395.

Kim, E., Sun, L., Gabel, C.V., and Fang-Yen, C. (2013). Long-Term Imaging of Caenorhabditis elegans Using Nanoparticle-Mediated Immobilization. PLOS ONE 8, e53419.

Larsen, P.L., Albert, P.S., and Riddle, D.L. (1995). Genes that regulate both development and longevity in Caenorhabditis elegans. Genetics 139, 1567–1583.

Manjarrez, J.R., and Mailler, R. (2020). Stress and timing associated with Caenorhabditis elegans immobilization methods. Heliyon 6, e04263.

Massie, M.R., Lapoczka, E.M., Boggs, K.D., Stine, K.E., and White, G.E. (2003). Exposure to the metabolic inhibitor sodium azide induces stress protein expression and thermotolerance in the nematode Caenorhabditis elegans. Cell Stress Chaperones 8, 1–7.

Murray, P., Hayward, S.A.L., Govan, G.G., Gracey, A.Y., and Cossins, A.R. (2007). An explicit test of the phospholipid saturation hypothesis of acquired cold tolerance in Caenorhabditis elegans. Proceedings of the National Academy of Sciences 104, 5489–5494.

Ohta, A., Ujisawa, T., Sonoda, S., and Kuhara, A. (2014). Light and pheromone-sensing neurons regulates cold habituation through insulin signalling in Caenorhabditis elegans. Nat Commun 5.

Podbilewicz, B., and Gruenbaum, Y. (2006). Live Imaging of Caenorhabditis elegans: preparation of samples. CSH Protoc 2006.

Rabin, Y., and Podbilewicz, B. (2000). Temperature-controlled microscopy for imaging living cells: apparatus, thermal analysis and temperature dependency of embryonic elongation in Caenorhabditis elegans. J Microsc-Oxf 199, 214–223.

Raizen, D.M., Zimmerman, J.E., Maycock, M.H., Ta, U.D., You, Y.J., Sundaram, M.V., and Pack, A.I. (2008). Lethargus is a Caenorhabditis elegans sleep-like state. Nature 451, 569–572.

Reeve, E.C.R. (1988). The Nematode Caenorhabditis elegans. Edited by William B. Wood and the Community of C. elegans Researchers. New York: Cold Spring Harbor Laboratory, 1988. 667 pages. US $97.00. ISBN 0 87969 307 X. Genetics Research 52, 243–244.

Robinson, J.D., and Powell, J.R. (2016). Long-term recovery from acute cold shock in Caenorhabditis elegans. BMC Cell Biol 17, 11.

Rohde, C.B., and Yanik, M.F. (2011). Subcellular in vivo time-lapse imaging and optical manipulation of Caenorhabditis elegans in standard multiwell plates. Nat Commun 2, 7.

Wang, Y.L., Grooms, N.W.F., Civale, S.C., and Chung, S.H. (2021). Confocal imaging capacity on a widefield microscope using a spatial light modulator. PLOS ONE 16, e0244034.

Ward, A., Liu, J., Feng, Z.Y., and Xu, X.Z.S. (2008). Light-sensitive neurons and channels mediate phototaxis in C-elegans. Nat Neurosci 11, 916–922.

Zeng, F., Rohde, C.B., and Yanik, M.F. (2008). Sub-cellular precision on-chip small-animal immobilization, multi-photon imaging and femtosecond-laser manipulation. Lab Chip 8, 653–656.

