## Supplementary material for "High-throughput submicron-resolution microscopy of entire *C. elegans* populations under strong immobilization by cooling cultivation plates": Figure S3

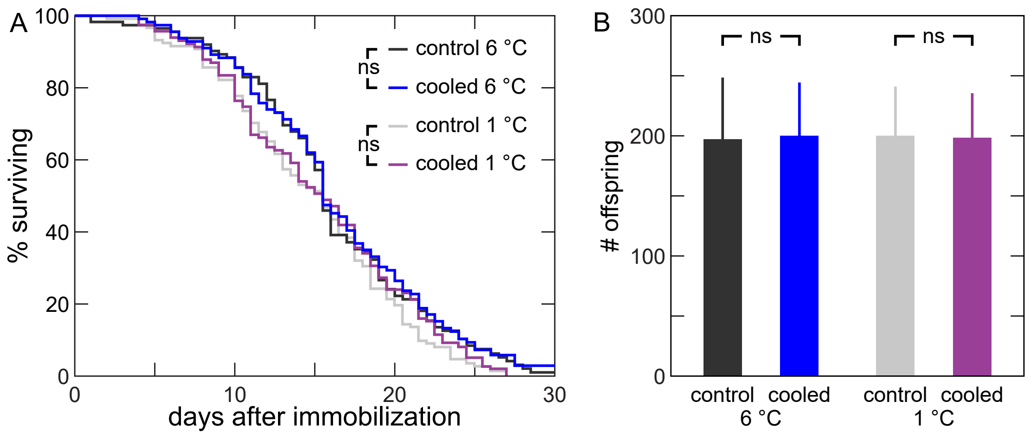


**Fig. S3. Cooling immobilization does not reduce *C. elegans* lifespan and fecundity.** Animal lifespan (a) and fecundity (b) after slow cooling to 6 °C or fast cooling to 1 °C, followed by 1 h at $T_{set}$. $n$ = 110 (lifespan), 55 (fecundity) per condition. $p < 0.05$ (*) and $p < 0.001$ (**).
