## Supplementary material for "High-throughput submicron-resolution microscopy of entire *C. elegans* populations under strong immobilization by cooling cultivation plates": Figure S2

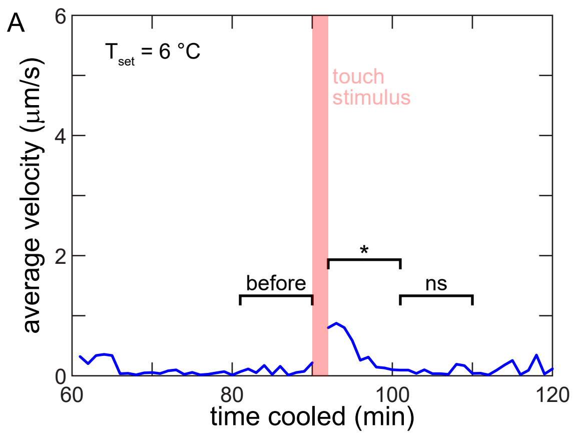


**Fig. S2. Touch stimulus briefly reactivates movement.** Nose tip velocity before and after harsh touch at 90 min are significantly different. Animals return to immobilization in 10 min. $n = 6$ animals. $p < 0.05$ (*).
