## Supplementary material for "High-throughput submicron-resolution microscopy of entire *C. elegans* populations under strong immobilization by cooling cultivation plates": Figure S1

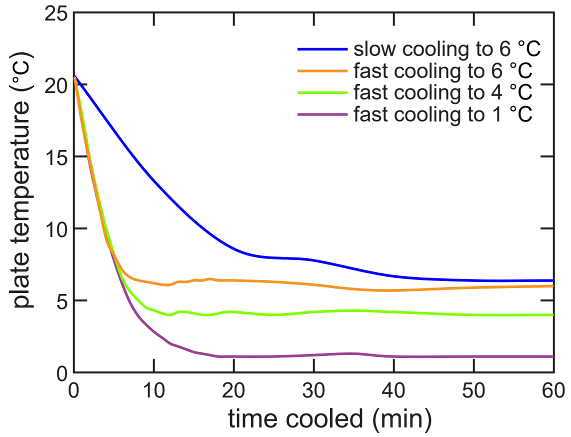


**Fig. S1. Plate cooling profiles.** Plate temperatures over time for different cooling methods and $T_{set}$.
