## Supplementary material for "High-throughput submicron-resolution microscopy of entire *C. elegans* populations under strong immobilization by cooling cultivation plates": Tab. S1

| class | examples | mechanism of action | drawbacks |
|---|---|---|---|
| chemical | sodium azide | block adenosine triphosphate production (Reeve, 1988; Bowler, *et al.*, 2006; Manjarrez and Mailler, 2020) | labor-intensive pre- and post-processing, stress-related secondary effects (Massie, *et al.*, 2003; Manjarrez and Mailler, 2020) |
| | levamisole | acetylcholine receptor overstimulation at neuromuscular junction (Brenner, 1974; Robertson, *et al.*, 2010; Puttachary, *et al.*, 2010; Martin, *et al.*, 2012) | labor-intensive pre- and post-processing, may impact viability and morphology, stress-related secondary effects, levamisole-resistant mutations (Martin, *et al.*, 2012) |
| mechanical | nanoparticle / microbead | pressure, friction (Kim, *et al.*, 2013) | requires slide mounting, small movements persist, can cause permanent damage, animals not accessible during immobilization (Kim, *et al.*, 2013) |
| | Pluronic F127 hydrogel (often used with microbead matrices) | viscosity, friction (Shachaf, *et al.*, 2010; Basak and Bandyopadhyay, 2013; Krajniak and Lu, 2010) | labor-intensive pre- and post-processing, forward-backward locomotion and other small movements persist (Krajniak and Lu, 2010; Krajniak, *et al.*, 2013; Dong, *et al.*, 2018) |
| | microfluidics, on-chip technologies | controlled flow, compression, suction (Chronis, *et al.*, 2007; Lockery, 2007; Zeng, *et al.*, 2008; Hulme, *et al.*, 2007; Guo, *et al.*, 2008; Ben-Yakar, *et al.*, 2009; Chronis, 2010; Crane, *et al.*, 2010) | complex setup, setup designed for specific applications, small movements persist (Chronis, *et al.*, 2007; Lockery, 2007; Ben-Yakar, *et al.*, 2009; Chronis, 2010) |
| extreme temperature | 2-4 °C cooling in multiwell or microfluidic | undefined cooling effects (Chung, *et al.*, 2008; Rohde and Yanik, 2011) | stress-related secondary effects, may cause damage to animal cells, small movements persist (Chung, *et al.*, 2008) |
| | 37 °C heating | light-induced heat knockdown (Chuang, *et al.*, 2013) | complex setup, stress-related secondary effects, high fatality under multiple immobilizations (Chuang, *et al.*, 2013) |

**Table S1.** Existing immobilization methods grouped by class and mechanism. Note that those techniques are often combined.
